# Phase variable expression of *pdcB*, a phosphodiesterase influences sporulation in *Clostridioides difficile*

**DOI:** 10.1101/2021.05.24.445537

**Authors:** Babita Adhikari Dhungel, Revathi Govind

**Affiliations:** Division of Biology, Kansas State University, Manhattan, KS, 66506, USA

**Keywords:** *Clostridioides difficile*, *C. difficile*, phase variation, gene regulation

## Abstract

*Clostridioides difficile* is the causative agent of antibiotic-associated diarrhea and is the leading cause of nosocomial infection in developed countries. An increasing number of *C. difficile* infections are attributed to hypervirulence strains that produce more toxins and spores. *C. difficile* spores are the major factor for the transmission and persistence of the organism. Previous studies have identified global regulators that influence sporulation in *C. difficile*. This study discovered that PdcB, a phosphodiesterase to influence sporulation in *C. difficile* UK1 strain positively. Through genetic and biochemical assays, we have shown that phase variable expression of *pdcB* results in hypo- and hyper-sporulation phenotype. In the “ON” orientation, the identified promotor is the right orientation to drive the expression of *pdcB*. Production of PdcB phosphodiesterase reduces the intracellular cyclic-di-GMP concentration, resulting in hyper-sporulation phenotype. The OFF orientation of *pdcB* switch or mutating *pdcB* results in increased cyclic-di-GMP and hypo-sporulating phenotype. Additionally, we demonstrated that CodY binds to the upstream region of *pdcB* to represses its expression, and CodY mediated repression is relieved by the DNA inversion.

## Introduction

Phase variation is a phenomenon where two distinct phenotypes are established within a clonal bacterial population in a reversible manner. This phenomenon in bacteria has been long recognized and has been well studied in various bacterial pathogens. Heterogeneity within a bacterial population is readily visible as colony variation and is often associated with pathogenicity (Jiang *et al*., 2019; Sekulovic *et al*., 2018; Silverman and Simon, 1980). Genes that are regulated by phase variation are often involved in pathogenesis, such as those that encode fimbriae, flagella, cell surface protein, capsular polysaccharides, components that are responsible for establishing an interaction with the host, which gives the bacteria an advantage against the challenging environment of host like, colonization, establishing an infection or evading from host immune response (Li and Zhang, 2019; Sekulovic *et al*., 2018). Phase variation has been reported in several Gram-negative bacterial pathogens, including *Salmonella* (Kazuhiro Kutsukake and Iino, 1980; K Kutsukake and Iino, 1980; Salehi *et al*., 2017; Scott and Simon, 1982; Silverman *et al*., 1979; Silverman and Simon, 1980; Smith and Selander, 1990), *E*.*coli* (Eisenstein, 1988; Klemm, 1986; McClain et al., 1993, 1991), *Klebsiella* (Struve *et al*., 2008), *Neisseria* (Tauseef et al., 2013; van der Ende *et al*., 2000) and *Campylobacter jejuni* (Bacon et al., 2001; Holst Sørensen *et al*., 2012) where it has been employed to evade host immune system, increase adhesion, colonization, and virulence. While there are ample reports on phase variable regions and their role in the pathogenesis of Gram-negative bacteria, little is known about the role of phase variation in Gram-positive pathogens. *Streptococcus pneumoniae* utilize phase variation to switch colony morphology and to express pilus (Li and Zhang, 2019). Phase variation has been reported more recently in *C. difficile*, where two distinct colony morphologies: smooth, circular, and rough filamentous, were observed and were found to be associated with motility and distinct cell shapes (Anjuwon-Foster and Tamayo, 2017; Garrett *et al*., 2019). In this study, we initiated the investigation to determine the molecular mechanisms involved in the phase variable sporulating (UK1_O) and non-sporulating (UK1_T) colonies observed in *C. difficile* UK1 strain.

*C. difficile* is an anaerobic pathogen that lives in the mammalian GI tract (Edwards and McBride, 2014). *C. difficile’s* growth in the GI tract is kept on check by the commensal gut microbiota. But during antibiotic treatment, a healthy bacterial population of the gut are disrupted thereby creating an environment that is favorable for *C. difficile* to proliferate and cause infection (Edwards and McBride, 2014). *C. difficile* infection is caused primarily by the toxin produced by the bacteria, which causes inflammation of gut epithelium leading to severe diarrhea, commonly known as antibiotic-associated diarrhea, and ultimately causing fetal pseudomembranous colitis Mullish and Williams, 2018; Prete *et al*., 2019). Each year >200,000 *C. difficile* infection (CDI) and >12,000 deaths due to CDI are reported in the United States (CDC, 2020). Due to the increased number of infections and a massive cost of health care of ∼ 4.2 billion dollars, CDI has been categorized by the CDC as an urgent threat to public health (Lessa *et al*., 2015). The increasing number of *C. difficile* infections has been partly attributed to the emergence of hypervirulent *C. difficile* strains, which produce more toxins and spores (Denève *et al*., 2009). *C. difficile* spores are the major cause of transmission and persistence and are the major factor for making it the number one cause of hospital-acquired infection in North America. Ingested spores make their way to the intestine where they germinate with the help of bile salts into active vegetative cells and secrete cytotoxins that cause tissue damage and inflammation (Paredes-Sabja *et al*., 2014). Since they are the root cause of infection, it is important to understand the regulation of sporulation in *C. difficile*. In this study, by characterizing the phase variation involving sporulating phenotypes, we have uncovered a new link between intracellular c-di-GMP concentrations and sporulation.

Cyclic-di-GMP is a common bacterial second messenger found in many bacterial species. Two classes of enzymes are important for c-di-GMP homeostasis in a bacterial cell, diguanylate cyclases (DGCs) that synthesize c-di-GMP from 2 GTP molecules and phosphodiesterases (PDEs) that degrade c-di-GMP. In *C. difficile*, c-di-GMP has been shown to negatively regulate toxin production (Bordeleau and Burrus, 2015; Hall and Lee, 2018; McKee *et al*., 2018b, 2013; Purcell *et al*., 2017a) and flagella mediated swimming motility (Bordeleau and Burrus, 2015; Hall and Lee, 2018; McKee *et al*., 2018b; Purcell *et al*., 2012), whereas positively regulate *pilA* mediated swarming motility (Bordeleau *et al*., 2015; Bordeleau and Burrus, 2015; Hall and Lee, 2018; McKee *et al*., 2018b, 2018a; Purcell *et al*., 2016), biofilm formation (McKee et al., 2018b; Purcell *et al*., 2017a), cell aggregation (Bordeleau *et al*., 2015; Purcell *et al*., 2012) and expression of cell wall anchor structures (Arato *et al*., 2019; Peltier *et al*., 2015). While c-di-GMP regulates a plethora of physiological processes, its influence on sporulation hasn’t been reported. In this study, we discovered the link between the reduced intracellular level of c-di-GMP and the hyper sporulation phenotype in *C. difficile* UK1 strain. Reduction in intracellular c-di-GMP level was brought about by the overexpression of *pdcB*, which encodes for a phosphodiesterase. We have also shown that *pdcB* mutant is hypo-sporulating and overexpression of the PdcB-EAL domain complements the sporulation phenotype. Our data shows that *pdcB* is expressed during the mid-exponential growth phase and is negatively regulated by CodY. When the *pdcB* gene is suppressed by CodY, it results in a non-sporulating colony. A DNA inversion in the upstream of *pdcB* gene partially relieves the repression by CodY, and expression of *pdcB* results in sporulation initiation in *C. difficile* UK1 strain.

## Results

### *C. difficile* UK1 strain exhibits phenotypic heterogeneity with two distinguishable colony morphologies

Earlier studies have shown that *C. difficile* strain R20291 belonging to ribotype 027, associated with epidemic infections, have phenotypic heterogeneity, thereby exhibiting two distinct colony morphologies *in vitro* (Anjuwon-Foster and Tamayo, 2017; Garrett *et al*., 2019). Consistent with these studies, we observed two distinct colony types in the UK1 strain during routine plating on TY agar (Fig. 1A). One colony type is round, has a circular and smooth edge, and is termed as an Opaque colony (UK1_O), while the other is irregular with a rough edge and is termed as Translucent colony (UK1_T). Under the phase-contrast microscope (Fig.1B), UK1_T cells were longer with an average length of 5.5 µm while the UK1_O cells are significantly shorter with an average length of 2.5 µm (Fig. 1C). Isolated UK1_T and UK1_O colonies were streaked on TY agar plates, and their morphologies were observed over time. Opaque colonies could maintain the colony morphology even after 24 h while the translucent colony gave rise to Opaque colonies after a day of incubation. Similar results were observed when they were grown in TY broth medium. Opaque colonies in the TY plate that were kept inside the anaerobic chamber for a prolonged time (>72 hrs) started exhibiting rough edges with the pale yellowish center over time, suggesting that opaque colonies could give rise to translucent colony type. Previous studies have also shown that streaking a culture from the center of the undifferentiated R20291 strain resulted in smooth colony type, while streaking from the edge resulted in rough colony types (Garrett *et al*., 2019).

**Figure 1.**
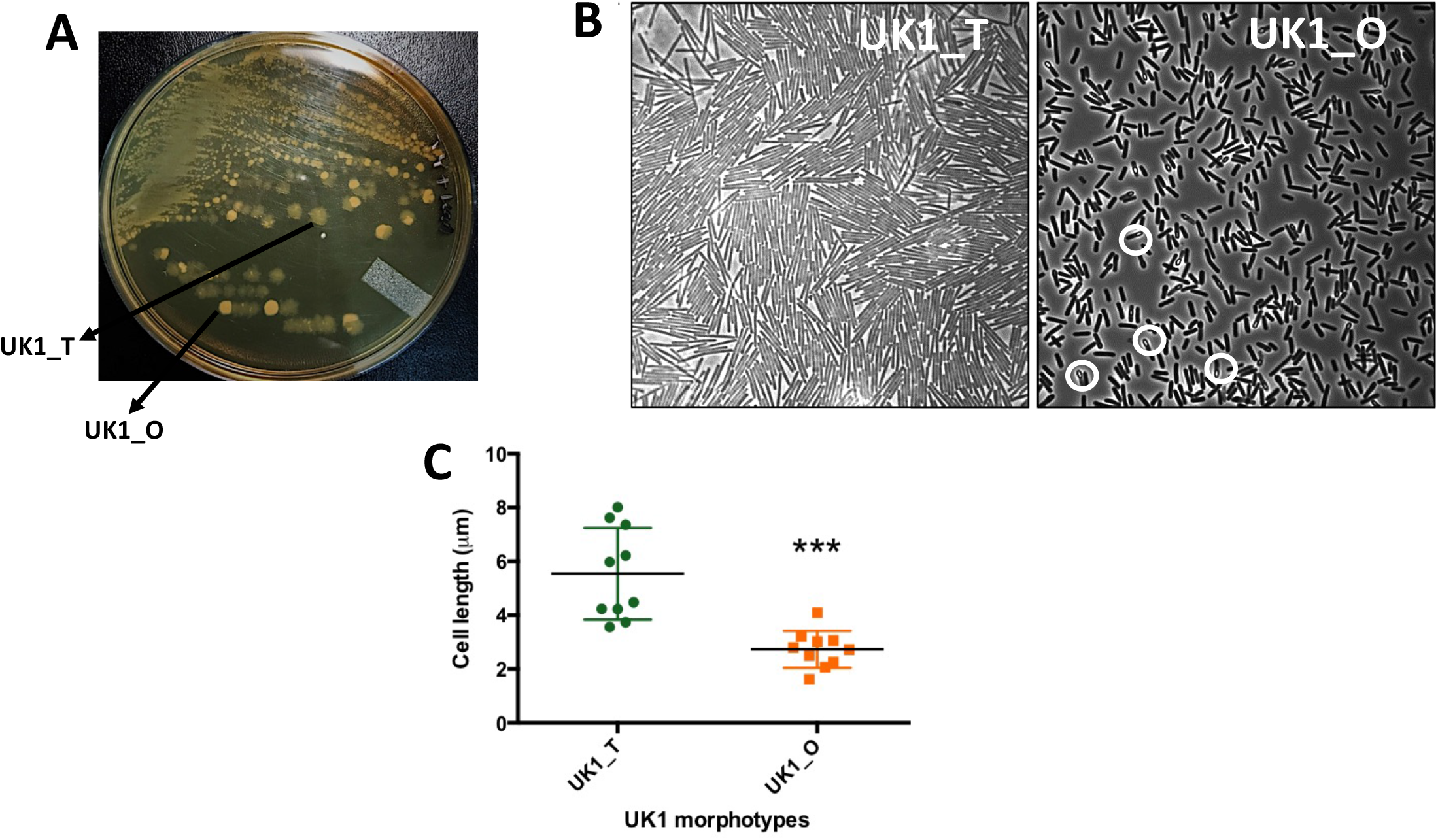
Colony and cell morphologies of UK1_T and UK1_O strains. **A**. UK1 strain, when grown in TY medium gives rise to phenotypically distinguishable Translucent (UK1_T) and Opaque (UK1_O) colony morphotypes. **B**. Phase-contrast microscopic images of UK1_T and UK1_O colony morphotypes. White circles highlight some of the sporulating cells in UK1_O. **C**. Average cell length showing UK1_T cells are significantly longer than UK1_O cells. Data analyzed using unpaired t-test with Welch’s correction where *** indicates P<0.0005.

### UK1_T and UK1_O colony morphologies exhibit distinct sporulation and swarming motility phenotype

In order to determine the physiological characteristics of these two colony types, we assayed for sporulation, toxin, and motility of UK1_T and UK1_O colony types. To determine the sporulation phenotype, we streaked the isolated UK1_T and UK1_O colonies in solid TY medium and we found that UK1_O strain was hyper-sporulating and produced significantly more spores compared to UK1_T strain at 24 hours (Fig. 1B, Fig. 2A). We also found that UK1_T and UK1_O strains produce similar level of cytoplasmic toxins at 16 h of growth (Fig. S1). While UK1_T and UK1_O strains exhibited similar swimming motility even after 72 hours in 0.5% agar TY medium (Fig. S1), we found that UK1_T strain exhibited enhanced swarming motility than the UK1_O strain when grown in TY medium with 1.8% agar at 72 hr (Fig. 2B). Similarly, UK1_T formed thick biofilm compared to UK1_O (Fig. 2C). Together, these data demonstrate that UK1_T and UK1_O are phenotypically different in sporulation, swarming motility, and biofilm formation. While the swarming motility and the biofilm formations were previously reported to be associated with phase variation in *C. difficile*, sporulation was not.

**Figure 2.**
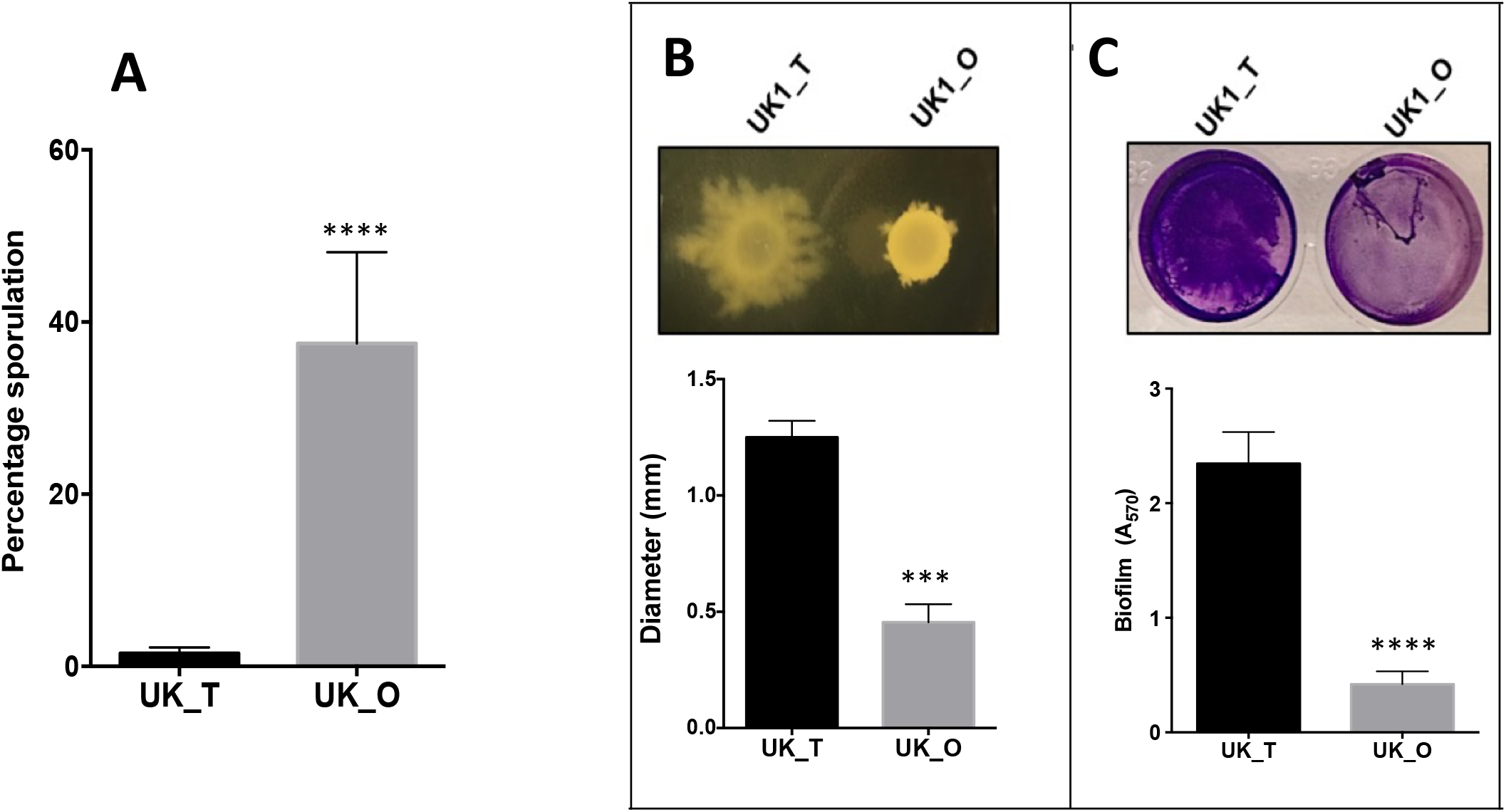
Phenotypic characterization of UK1_T and UK1_O strains. **A**. Percentage sporulation of UK1_T and UK1_O strains showing hyper sporulation phenotype in UK1_O morphotype. **B**. Swarming motility exhibited by UK1_T and UK1_O in TY+1.8% agar. **C**. Biofilm formation of UK1_T and UK1_O morphotypes. OD_570_ nm was measured to quantify the biofilm formation. Three biological replicates were used to carry out the experiments, and data were analyzed by two-tailed unpaired t-test with Welch’s correction where **** =P<0.00005. ***=P<0.005

### UK1_O strain undergoes phase variation by DNA inversion

To determine the genetic basis for the phenotypic heterogeneity that gave rise to translucent and opaque colonies in the UK1 strain, we extracted genomic DNA from UK1_T and UK1_O colony morphotypes and performed whole-genome sequencing at Tufts University Genomic core facility. The Sequenced genome was assembled against the closely related and annotated R20291 genome (Stabler *et al*., 2009). Unlike our expectation, the sequencing result did not show mutations in any of the known sporulation-associated genes. However, the intergenic region between *CDR20291_0684* and *CDR20291_0685* genes was listed among unassigned new junction evidence.

In order to determine if there is any information in this intergenic region, we amplified and cloned ∼1.5 kb intergenic region from UK1_T and UK1_O in a vector and sent 10 clones from each for sequencing. We aligned the sequences of UK1_T and UK_O to the R20291 genome (Stabler *et al*., 2009) and analyzed with NCBI BLASTn. The result suggested that all clones from UK1_T perfectly aligned with the published sequence of the R20291. In contrast, all the clones from UK1_O had a mismatch in the ∼200 bp segment of the intergenic region (Fig. S2), showing DNA inversion in the region.

### DNA inversion regulates the expression of the downstream gene *CDR20291_0685*

DNA inversion is known to regulate the expression of the downstream gene as it gives rise to two orientation of the loci where “ON” orientation enhances and “OFF” orientation represses the expression of the downstream gene or operon (Jiang *et al*., 2019; Silverman and Simon, 1980; Van de Putte and Goosen, 1992; Zieg and Simon, 1980). We performed a reporter fusion assay to determine the effect of DNA inversion in the expression of the downstream CDR20291_0685 gene. We fused ∼1.5 kb of the upstream DNA segment of *CDR20291_0685* containing the invertible region from both UK1_T and UK1_O strain with *gusA* reporter gene coding beta-glucuronidase to create plasmid constructs pBA042 and pBA045 (Table S1). Using site-directed mutagenesis, we mutated the inverted repeat sequences that flanked the invertible region to lock the upstream region either in the translucent (pBA043) or opaque (pBA046) orientation. The plasmid constructs were then introduced into the UK1 undifferentiated parent strain by conjugation. The strains were grown anaerobically in TY broth medium at 37°C and were harvested at different time points to perform the reporter gene expression assay. We observed significantly higher beta-glucuronidase activity at each time point with the reporter fusion that was locked in opaque orientation (Fig. 3A). Very minimal reporter activity was recorded with the reporter fusion that was locked in the translucent orientation. This observation is consistent with our qRT-PCR results, where we detected elevated levels of *CDR20291_0685* transcript in UK1_O when compared to that from UK1_T (Fig. 3B). Taken together, these results suggest that DNA inversion in the upstream region of the *CDR20291_0685* gene resulted in upregulation of its expression in the opaque UK1_O colonies.

**Figure 3.**
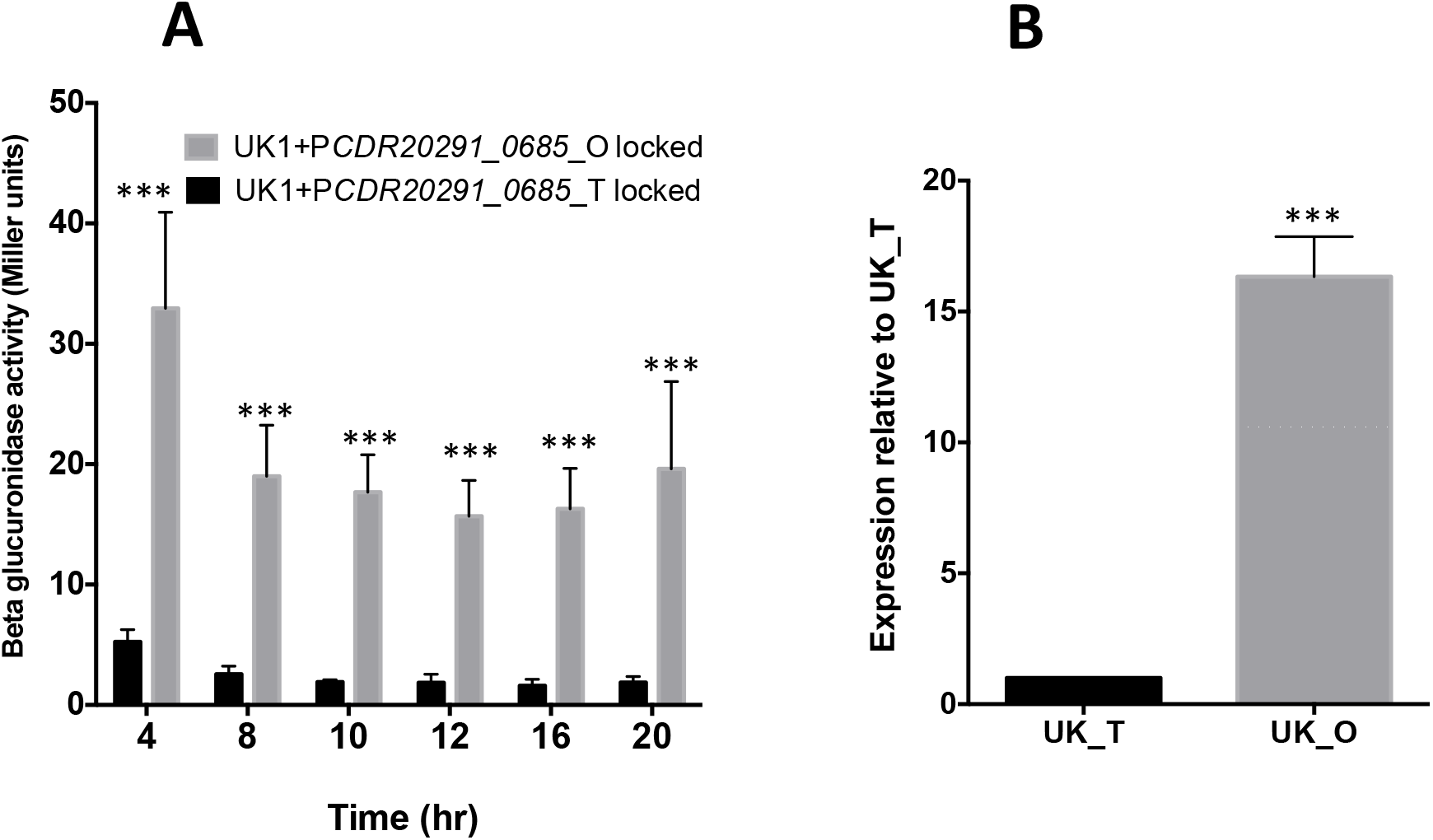
DNA inversion upregulates the expression of *CDR20291_0685* in UK_O strain. **A**. The Beta-glucuronidase activity of the P*CDR20291_0685* upstream*-gusA* fusions locked in UK1_T and UK1_O orientation in the UK1 parent strain. The data represents the results from 3 biological replicates. Data were analyzed using 2-way ANOVA (Sidak’s multiple comparisons test) comparing the mean of UK1_T and UK1_O at each time point. *** indicates P<0.0005. **B**. qRT-PCR results showing overexpression of *CDR20291_0685* in UK1_O strain at 16-hour time point. The representative results from three independent experiments are shown. Data were analyzed by a two-tailed unpaired t-test with Welch’s correction where ***=P<0.0005

### *CDR20291_0685* promoter lies within the invertible region

The mechanism by which DNA inversion regulates the expression of the gene depends upon its location. If DNA inversion occurs in the region that contains the promoter, it will disrupt the promoter, and the gene is not expressed (Eisenstein, 1988; Zieg and Simon, 1980). If the promoter is upstream of the switch, then an intrinsic terminator can be formed to terminate the transcription of the gene (Sekulovic *et al*., 2018). To determine the mechanism by which this inversion regulates the gene expression, we sought to identify the promoter of *CDR20291_0685*. We hypothesized that the promoter of *CDR20291_0685* is within the invertible region. We extracted total RNA from a 4 h culture of UK1_O strain and synthesized *CDR20291_0685* specific cDNA. We carried out PCR using this cDNA as a template and several primers spanning the inverted repeat sequence (Fig. 4A). Only the primer pairs ORG921/ORG853, and ORG922/ORG853 gave amplification product which suggests that the promoter region is located around the region spanned by the primer ORG921 (Fig. 4B). Consistent with this observation, we mapped the transcription start of *CDR20291_0685* using 5’RACE. Transcription is initiated 874-bp upstream of the *CDR20291_0685* translational initiation codon and is located next to the right repeat sequence (Fig. 4A). Hexamers -10 (TATTT) and -35 (TTGATA) separated by an -17-bp spacer are found upstream of the transcriptional start site.

**Figure 4.**
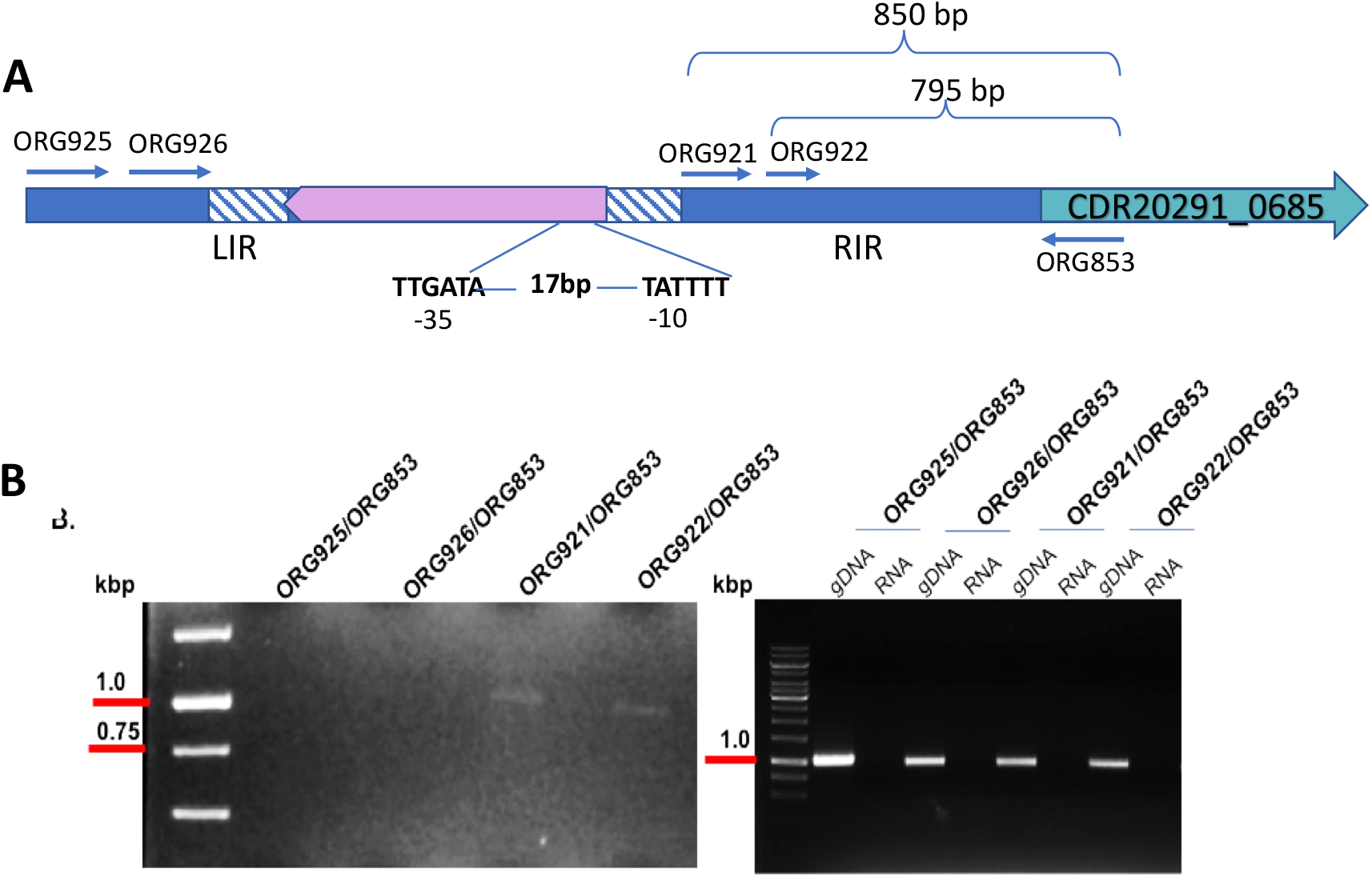
The promoter of *CDR20291_0685* is located within the invertible region. **A**. Schematic of the upstream region of *CDR20291_0685* from UK1_O strain (the orientation in which *CDR20291_0685* is expressed) depicting the invertible region flanked by the right and left inverted repeats (RIR and LIR). The forward and the reverse primers designed along the length of the invertible region are shown. The transcription initiation site was found near the RIR, and - 35 and -10 sites are marked. **B**. RT PCR detecting the transcripts of *CDR20291_0685*. Genomic DNA (gDNA) and RNA were used as positive and negative controls, respectively. Only the primer pairs ORG921/ORG853, and ORG922/ORG853 gave amplification products of size 882 bp and 825 bp, respectively.

### Overexpression of *CDR20291_0685* results in reduced c-di-GMP levels in UK1

*CDR20291_0685* is the orthologous gene of *CD630_07570* in strain CD630. *CD630_07570* hydrolyzes c-di-GMP to pGpG *in vitro* and therefore is a phosphodiesterase enzyme (Bordeleau *et al*., 2011). Therefore, we named *CDR20291_0685* as *pdcB* (<u>P</u>hospho<u>d</u>iesterase of *<u>C</u>. difficile* <u>B</u>). We have shown that DNA inversion in UK1_O strain results in the expression of *pdcB* (Fig. 3AB). To determine if overexpression of *pdcB* alters the levels of intracellular c-di-GMP in *C. difficile*, we first created UK1::*pdcB* mutant strain using ClosTron mutagenesis (Fig. S3) and extracted and measured the levels of intracellular c-di-GMP from UK1_T, UK1_O and UK1::*pdcB* strains. Our data suggest that UK1_T and UK1::*pdcB* strains have higher levels of c-di-GMP while UK1_O strain has significantly reduced c-di-GMP level (Fig. 5A). This result indicates that DNA inversion overexpresses the *pdcB* gene, which cleaves c-di-GMP and reduces the global intracellular concentration of c-di-GMP in the UK1_O strain.

**Fig 5.**
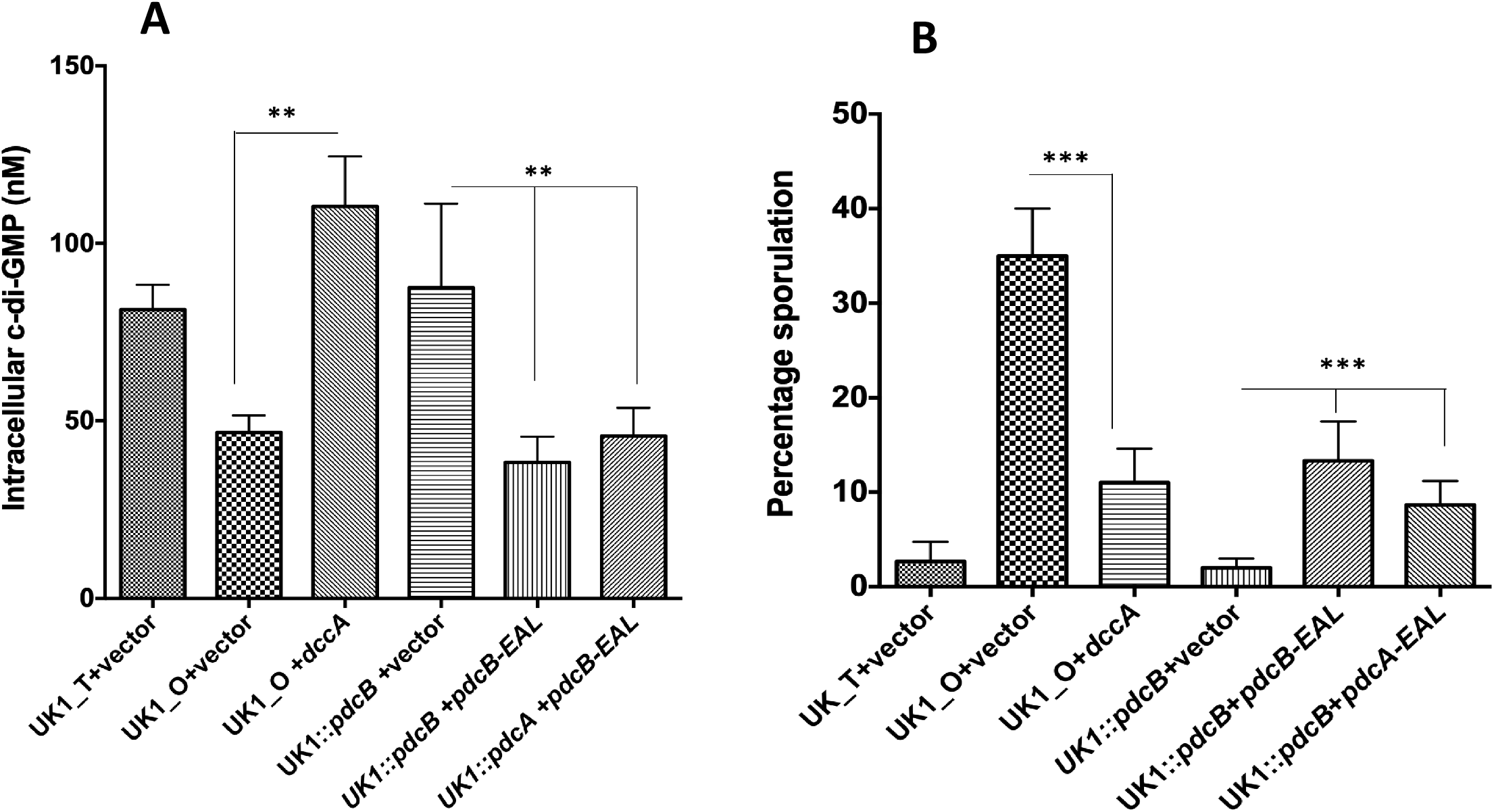
Intracellular c-di-GMP levels influence sporulation in UK1 strain. **A**. The intracellular levels of c-di-GMP (A) and sporulation percentage (B) in strains UK1_T, UK1_O, UK1_O expressing *dccA*, UK1::*pdcB* mutant, and its complemented strains. Three biological replicates were used to carry out the experiment, and data were analyzed by one-way analysis of variance (ANOVA) where ** indicates P<0.005, *** is P<0.0005.

### Sporulation in *C. difficile* UK1 is associated with the intracellular levels of c-di-GMP

Intracellular c-di-GMP homeostasis is maintained by two classes of enzymes, diguanylate cyclases that synthesize c-di-GMP and phosphodiesterases that hydrolyze c-di-GMP. Phosphodiesterases consist of either the EAL domain or an inactive GGDEF and active EAL domain (Hall and Lee, 2018). PdcB phosphodiesterase contains and both the GGDEF and EAL domains, and the *in vitro* assays have shown PdcB can cleave c-di-GMP (Bordeleau *et al*., 2011). Our results so far showed that DNA inversion upstream of *pdcB* regulated its expression and hence controlled the intracellular levels of c-di-GMP in *C. difficile* UK1, which in turn resulted in two phenotypic variants UK1_T, and UK1_O. We also observed a significant difference in the rate of sporulation between UK1_T, UK1_O. To determine if the intracellular levels of c-di-GMP is associated with a sporulation phenotype, we quantified percentage sporulation in UK1_T, UK1_O, UK1::*pdcB* mutant, and UK1::*pdcB* mutant complemented with plasmid-encoded PdcB-EAL domain under a tetracycline-inducible promoter. Despite multiple attempts, we were not able to complement the UK1::*pdcB* mutant strain with a full-length plasmid-encoded *pdcB*. Other studies have shown that complementation by just the EAL domain with the phosphodiesterase activity of *pdcA* is sufficient to rescue the associated motility phenotypes (Purcell *et al*., 2017b). Since our objective was to determine the role of intracellular levels of c-di-GMP in regulating sporulation in *C. difficile* UK1 strain, we performed the sporulation assays with the mutant strain complemented with the plasmid that encodes just the PdcB-EAL domain. Percentage sporulation data showed hypo-sporulating phenotype in UK1::*pdcB* strain was partly complemented when the PdcB-EAL was produced in the mutant strain (Fig. 5B). To further confirm that sporulation phenotype is associated with intracellular levels of c-di-GMP, we complemented the UK1::*pdcB* mutant with the plasmid-borne PdcA-EAL domain. PdcA is another well-characterized phosphodiesterase of *C. difficile*, and the PdcB shares homology in the conserved GGDEF and EAL domain with PdcA-EAL. We observed that sporulation phenotype was complemented with the PdcA-EAL domain as well. Additionally, we expressed *dccA* coding for the diguanylate cyclase from the *tet* inducible promoter to artificially increase the concentration of c-di-GMP and monitored the sporulation percentage upon the induction of *dccA*. As expected, the production of DccA significantly reduced the sporulation in the UK1_O strain (Fig. 5B). Measuring c-di-GMP in strains expressing *pdcA-EAL* or *pdcB-EAL* or *dccA* further confirmed their effect on c-di-GMP concentration (Fig. 5A).

The intracellular level of c-di-GMP is known to positively regulate swarming motility in *C. difficile* (Bordeleau *et al*., 2015). Consistent with previous studies, we observed increased swarming motility in the UK1_T strain, which has increased intracellular c-di-GMP levels compared with the UK_O strain. So, to further confirm that complementation of the UK1::*pdcB* mutant with PdcB-EAL and PdcA-EAL domains reduces the intracellular c-di-GMP and sporulation phenotype that we observed is associated with the intracellular c-di-GMP levels, we carried out a swarming motility test assay for all the strains (Fig. S4). As expected, we observed decreased swarming motility in the complemented strains similar to the UK1_O strain, which has reduced intracellular c-di-GMP levels. Taken together, these data suggest that the intracellular levels of c-di-GMP influence sporulation in *C. difficile*.

### CodY binds to the upstream region and represses the expression of *pdcB*

Several studies have shown CodY mediated regulation of c-di-GMP signaling proteins. Microarray analysis in the CD630 strain has identified two cyclic di-GMP signaling proteins, *CD630_07570* (*pdcB*) and *CD1476*, among the 140 genes that were regulated by CodY (Dineen *et al*., 2010). The study demonstrated enhanced expression of both of these proteins in the *codY* mutant and identified a consensus CodY binding site in the upstream regions of *CD1476* and *CD630_07570 (pdcB)*. Another study has shown that CodY binds directly to the promoter region and represses the expression of *pdcA*, a phosphodiesterase that cleaves c-di-GMP in the JIR8094 strain (Purcell *et al*., 2017b). To determine if the expression of *pdcB* is controlled by CodY, we first determined whether a CodY binding consensus sequence is present in the upstream region of *pdcB*. The two potential CodY binding sites upstream of *pdcB* are TTTTTAGAAAAGTA and AAATTTTCAATGAAT within the region that undergoes inversion (Fig. 6A). CodY binding site in the opaque orientation is upstream of the predicted promoter region in the UK1_O orientation, which can explain the higher expression of *pdcA* in UK1_O when compared to UK1_T. We hypothesized that CodY binding would completely shut down even the low-level expression of *pdcB* in the UK1_T strain. DNA inversion in the UK1_O strain would shift the CodY binding site to the upstream region of the predicted promoter, leading to the expression of *pdcB*.

**Figure 6.**
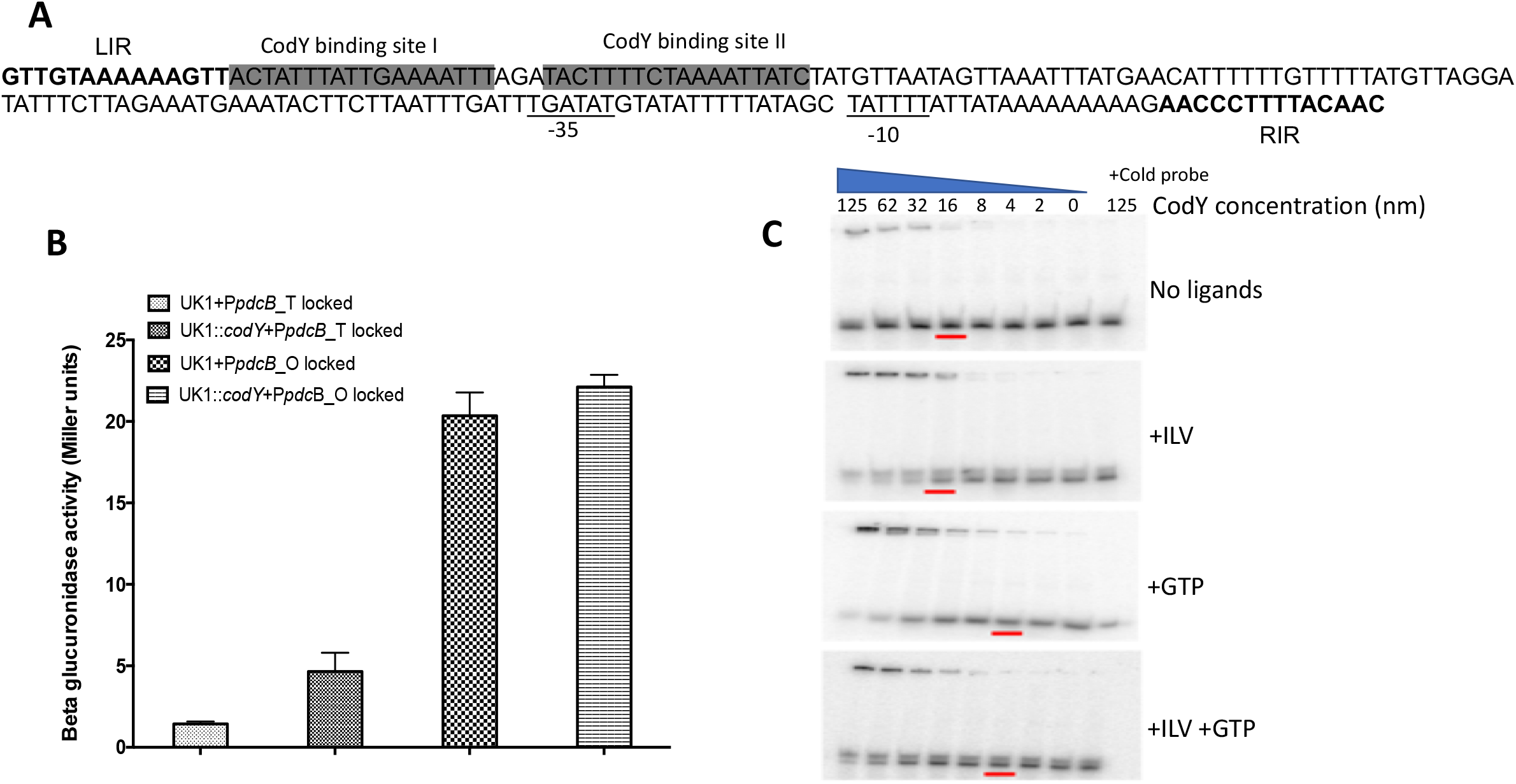
CodY represses the expression of *pdcB*, which is partially relieved by DNA inversion. **A**. Schematic showing the potential CodY binding sites that lie within the invertible region in upstream of *pdcB*. **B**. Beta-glucuronidase activity of P*pdcB-gusA* fusions locked in either translucent or opaque orientation in the parent UK1 and UK1::*codY* mutant. The data shown are the standard errors of the mean of three biological replicates. Statistical analysis was performed using two-way ANOVA.**=P<0.05.**C**. Binding of purified CodY to the upstream region of *pdcB*. Binding was increased with increasing concentration of CodY. The presence of GTP and ILV further enhanced CodY binding. Ten-fold excess of the cold probe was added with 125 ng of purified CodY in a control reaction to rule out non-specific binding.

To determine if CodY regulates the expression of *pdcB*, we carried out the reporter fusion assay. We used the same reporter fusions (pBA040, pBA043, pBA046) that were used earlier in this study. Each of the constructs was introduced into the UK1 undifferentiated parent strain and UK1::*codY* mutant strain by conjugation. The strains were grown anaerobically in TY broth medium at 37°C, and cells were harvested at different time points to perform the reporter assay. We observed increased beta-glucuronidase activity at each time point when it was expressed from the *pdcB* upstream locked in opaque orientation, with higher expression at mid-exponential phase from 4-12 hours for both UK1 parent and UK1::*codY* mutant strains (Fig. 6B). Minimal reporter activity was observed with the upstream locked in translucent orientation. However, in the absence of CodY, an approximately 1.5-fold increase in the expression could be seen from the promoter locked in the translucent orientation, suggesting the presence of a second but much weaker promoter upstream of the CodY binding site.

To further confirm the interaction of CodY with *pdcB* upstream DNA, we carried out Electrophoretic Mobility Shift Assay (EMSA) with purified CodY protein. We synthesized the DNA fragment of 47bp containing the potential CodY binding site and radio-labeled with γ-^32^ P and carried out the binding of the fragment with purified CodY protein with increasing concentration of the protein, both in the absence and presence of its cofactors ILV (Branched-chain amino acids Isoleucine, Leucine, and Valine) and GTP (Fig. 6C). The shift in the bands at higher concentrations of CodY protein suggests that purified CodY binds to the DNA fragment, and the binding was enhanced in the presence of its cofactors GTP and ILVs. These data suggest that CodY has a direct role in downregulating the expression of *pdcB*. Taken together, these results suggest that *pdcB* expression in UK1 is tightly regulated by both DNA inversion and by the activity of CodY.

## Discussion

The major objective of this study was to determine the role of phase variation in the physiology of *C. difficile* UK1 strain. The genome-wide analysis identified seven different types of invertible sites in the *C. difficile* genome, designated as Cdi1 to Cdi7 (Sekulovic *et a*l., 2018). Previous studies have shown that Cdi2 to present in the upstream region of *pcdB* (*CDR20291_0685*), encoding one of the c-di-GMP phosphodiesterases (PDEs) in *C. difficile* UK1 strain (Sekulovic *et al*., 2018). In this work, we have shown the role of Cdi2 in regulating the expression of *pdcB*. Our data show that DNA inversion in the upstream of *pdcB* in UK1_O strain overexpresses *pdcB*, and this genotype is associated with reduced intracellular c-di-GMP. Consistent with this result, the UK1_T strain, which had minimal *pdcB* expression and UK1::*pdcB* mutant, demonstrated relatively higher levels of c-di-GMP. *C. difficile* is predicted to have 18 predicted PDEs (Bordeleau *et al*., 2011). Despite the redundancy, intracellular c-di-GMP levels seem to be greatly affected by the expression of *pdcB*.

While c-di-GMP regulates a plethora of physiological processes in *C. difficile*, its role in sporulation had not been previously described. Only a few studies have investigated the role of c-di-GMP in sporulation. Recently, the relationship between c-di-GMP and sporulation has been explored in *Streptomyces venezuelae*, a Gram-positive soil bacterium, where low levels of c-di-GMP resulted in hyper sporulation phenotype (Al-Bassam *et al*., 2018; Gallagher *et al*., 2020; Hengst *et al*., 2010; Latoscha *et al*., 2019; Liu *et al*., 2019; Tschowri *et al*., 2014). Gallagher KA, *et al*. reported that c-di-GMP bridges the binding of anti-sigma factor RsiG with sigma factor WhiG, strengthening their interaction and preventing sigma factor WhiG-dependent transcription of late sporulation genes (Gallagher *et al*., 2020). Similarly, another study demonstrated that c-di-GMP is required to control the transition from vegetative to reproductive cells (Bush *et al*., 2015). Their study reported that c-di-GMP was needed to form the active dimeric form of BldB, the master regulator of cell development, and represses the development of reproductive hyphae and keeps *Streptomyces* in their vegetative state (Al-Bassam et al., 2018; Bush et al., 2015; Gallagher et al., 2020; Hengst et al., 2010). These studies suggest that high c-di-GMP level traps *Streptomyces* in vegetative growth, and low c-di-GMP levels cause precocious hyper sporulation. Single-cell microscopy studies carried out in *B. subtilis* have shown a positive correlation of high c-di-GMP levels with sporulation (Weiss et al., 2019). Our current work is the first study to associate intracellular c-di-GMP concentration to sporulation in *C. difficile* directly. Hyper sporulation was observed when the intracellular c-di-GMP concentration was reduced by ∼1.5 fold in UK1_O strains than the UK1_T strains grown *in vitro*. It is noteworthy that, in our previous study, R20291::*sinRR’* mutant strain, which is asporogenic, had elevated levels of c-di-GMP compared with the R20291 parent strain (Girinathan et al., 2018). This study further corroborates our current observation. By analyzing percentage sporulation in UK1::*pdcB* mutant strain and PdcB-EAL and PdcA-EAL domains complemented strains, we have shown that production of phosphodiesterase that reduces c-di-GMP concentration is positively associated with sporulation phenotype. However, the overexpression of EAL domains of both PdcB and PdcA have only partially complemented the sporulation phenotype. This could be because of the overexpression of the PdcB-EAL domain alone without the associated GGDEF domain. Although the GGDEF domain of PdcB-EAL is catalytically inactive (Sekulovic et al., 2018), it might be necessary to enhance the activity of the EAL domain and fully complement sporulation. In homologous proteins which contain both GGDEF and EAL domains, catalytically inactive GGDEF domains are capable of binding to GTP, thus enhancing the PDE activity of the neighboring EAL domain (Christen et al., 2005).

Regulation of *C. difficile* sporulation can occur either at the transcriptional level by altering the expression of *spo0A* or at the post-translational level by interfering with the phosphorylation of Spo0A (Edwards and McBride, 2014). There are two known mechanisms by which c-di-GMP is known to function. First, c-di-GMP mediated regulation can occur through RNA-based effectors, which includes direct binding of c-di-GMP to riboswitches present in the 5’ UTR of the target gene transcript leading to premature termination of the transcript. Second, c-di-GMP mediated regulation can occur through protein effectors containing PilZ domain, GMP binding domain, diguanylate cyclases containing I-sites, GIL proteins, MshEN domains, and the Cle subfamily of CheY proteins which sense and respond to the changes in the intracellular c-di-GMP concentration (McKee *et al*., 2018b). Whether c-di-GMP binding riboswitches or c-di-GMP binding domains are present in the histidine kinases that phosphorylate Spo0A, and whether c-di-GMP regulates their expression or activity, thus regulating sporulation needs to be further investigated. On the other hand, c-di-GMP could indirectly influence sporulation by affecting the expression of regulators known to affect *spo0A*.

Previous studies have shown that *C. difficile* UK1::*codY* mutant exhibits a hyper-sporulation phenotype suggesting that CodY represses sporulation (Nawrocki *et al*., 2016). The exact mechanism by which CodY regulates sporulation is not understood. However, studies have shown that CodY directly regulates the expression of *sinRI* (Girinathan *et al*., 2018) and *opp* and *app* (Edwards *et al*., 2014; Nawrocki *et al*., 2016), which are positive regulators of sporulation. Results in this study suggest that phosphodiesterases can play a role in regulating sporulation. This could be indirectly by controlling CodY activity, which is directly regulated by intracellular GTP concentration. It has been shown that increasing the concentration of GTP enhances the phosphodiesterase activity of PdcA, suggesting the role of intracellular GTP in PDE activity (Purcell *et al*., 2017b) and might be important for PdcB activity as well. In the presence of GTP, CodY directly binds to upstream regions of *pdcB* and *pdcA* and represses their expression, reducing the degradation of GTP into c-di-GMP. However, any PdcB/PdcA produced would be competing for GTP with CodY, and activation of their enzymatic activity in the presence of GTP would reduce the GTP concentration and, in turn, CodY activity. Thus active CodY can repress the expression of pdcA/pdcB, and active PdcA/PdcB can influence CodY activity. This regulatory loop could be important to control the sporulation initiation and other CodY regulated pathways. It is however, important to note that DNA inversion plays a significant role in *pdcB* expression than CodY.

While we were working on characterizing *pdcB* role in UK1 phase variation, another study reported the role of *cmrRST* on phase variation in *C. difficile* R20291 strain (Garrett *et al*., 2019). Similar to the *pdcB* gene, the *cmrRST* operon carries a DNA inversion element belonging to *cdi6* group. CmrR and CmrS is a two-component signal transduction system coding for a response regulator and a histidine kinase, respectively. To check any connection between *cmrRST* and phase variation in the UK1 strain, we checked the orientation of the *cmrRST* upstream region in the UK1_O and UK1_T strains. In the UK1_T colonies, the PCR product indicating the OFF orientation was in higher concentration compared to the product indicating ON orientation (Fig. S5). In the UK1_O colonies, the *cmrRST* operon was observed to be in the ON orientation. In the R20291 strain, the rough colonies (similar to UK1_T) were produced when the *cmrRST* operon was in the ON orientation, and smooth colonies (similar to UK1_O) resulted when the *cmrRST* was in OFF orientation. In our results, we observed the opposite. Further investigation is needed to check whether *cmrRST* has any role in phase variation in the UK1 strain.

In Summary, our study has shown phase variation mediated regulation of intracellular c-di-GMP homeostasis and its effect on *C. difficile* physiology (Fig. 7). Although both translucent and opaque strains are shown to produce similar levels of toxins, they differ drastically in other phenotypes like sporulation and colony morphology. Based on the swarming motility and biofilm formation phenotypes, it can be suggested that during initiation of infection, UK1 might prefer to exist as the translucent colony cells so that they can better colonize the host and cause infection. Under the selective pressure exerted in the intestinal tract and host immune responses, the UK1 strain might prefer to switch to opaque colony type and activate the signaling cascade to produce more spores so that it can evade the host immune response and persist in the host to cause recurrent infection or in the environment once it is released outside of the infected individual. Understanding the importance of phase variable expression of *pdcB* and its associated phenotypes during *in vivo* infection model is a priority for future investigation and will shed light on the importance of *pdcB* in *C. difficile* pathogenesis.

**Figure 7.**
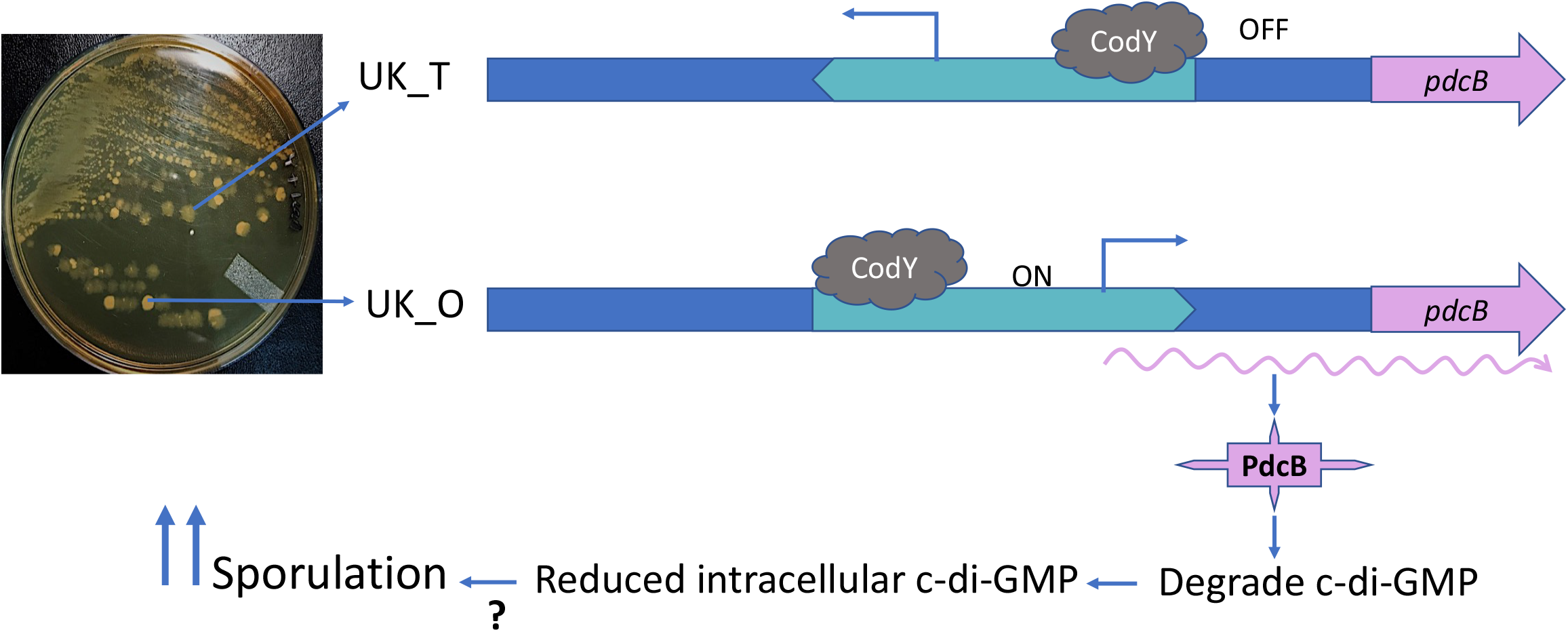
Proposed model. In UK_T cells, the predicted promoter region of *pdcB* is in the inverted orientation. The binding of CodY downstream further blocks any transcription of the *pdcB*. In the UK_O opaque cells, because of the DNA inversion, the predicted promoter is in the right orientation and drives the expression of *pdcB*, and is also not affected by the CodY binding to the upstream. Increased production of PdcB results in decreased c-di-GMP, which in turn influences the sporulation rate.

## Material and Methods

### Bacterial strains and growth conditions

*C. difficile* UK1 strains (Table S1) were grown in TY (Tryptose and Yeast extract) medium agar or broth culture in an anaerobic chamber maintained at 10% H_2_, 10% CO_2,_ and 80% N_2_. Transconjugants were selected and grown in TY agar with lincomycin (Linc 20µg/ml) or thiamphenicol (Thio; 15 µg/ml) or both. An *E*.*coli* strain optimized for conjugation S17-1 was used for conjugation (Teng *et al*., 1998), and the *E*.*coli* DH5α strain was used for cloning. *E*.*coli* strains were cultured aerobically in LB (Luria-Bertani) supplemented with ampicillin (100 µg/ml) or chloramphenicol (25 µg/ml) when necessary.

### Phase-contrast microscopy

*C. difficile* strains were grown in TY medium as described above. One ml of overnight culture was centrifuged at 17,000g for 1 min and washed with 30μl of sterile PBS. The resulting pellets were fixed for 2 hours at room temperature using 2% paraformaldehyde in 1X PBS. A thin layer of 0.7% agarose was applied to the surface of a microscopic slide, and 2μl of concentrated culture was placed on it. The cells were imaged using Zeiss LSM-5 PASCAL (objective lens 100x/1.4 oil). The LSM 5 service pack was used to acquire the images of at least three fields for each strain.

### Sporulation assay

Sporulation assays were performed in TY medium as described previously (Girinathan *et al*., 2017). To induce germination of any spores in the inoculum, *C. difficile* cultures were grown overnight in TY broth supplemented with 0.1% taurocholate. Cells were then diluted in TY medium to an OD600 of 0.5, and 100 μl the OD600 adjusted culture was spread on TY agar. Plates were incubated at 37°C for 36 hours. To enumerate the number of viable spores, cells were harvested using 1 ml of TY broth, and 500μl of the samples from each culture were mixed 1:1 with 95% ethanol and incubated for 30 minutes at room temperature to kill all the vegetative cells. The ethanol-treated samples and the untreated samples were then serially diluted, plated on TY agar with 0.1% taurocholate, and incubated at 37°C for 24 to 48 hours. The percentage of ethanol-resistant spores was calculated by dividing the number of CFU from spores by the total number of CFU and multiplying the value by 100. The results were based on a minimum of three biological replicates.

### RNA isolation, Reverse Transcriptase (RT-PCR) and Quantitative reverse transcriptase PCR (qRT-PCR)

*C. difficile* cultures were grown in TY medium, and 6 ml of cells were harvested at different time points by centrifugation at 3000g for 30 min at 4 °C. Total RNA was extracted from the harvested cells following the protocol described previously (El Meouche *et al*., 2013). Total RNA was treated with DNase (Turbo; Ambion) for 2 hours at 37 °C. The cDNA was synthesized using 5μg of total RNA at 42 °C for 2 hours using avian myeloblastosis virus (AMV) reverse transcriptase (Promega). A 20 µL reaction containing 10 ng or 10 pg (for 16S rRNA) of cDNA, 400 nM gene-specific primers, and 12.75 µL of SYBR PCR master mix (BioRad) was used to perform Real-time quantitative PCR using iQPCR real-time PCR instrument (BioRad). Amplification and detection were performed as described previously (El Meouche *et al*., 2013). The quantity of cDNA of a gene in each sample was normalized to the quantity of *C. difficile* 16S rRNA gene, and the ratio of normalized target concentrations (threshold cycle [2^-ΔΔCt^] method) (El Meouche *et al*., 2013; Saujet *et al*., 2011) gives the relative change in gene expression. The cDNA prepared was also used as a template to perform reverse transcriptase PCR to detect *pdcB* transcripts using forward primers ORG925, ORG926, ORG921, ORG922, along with the reverse primer ORG 853 (Table S2).

### 5’RACE

5′RACE (Rapid amplification of cDNA ends) assays were performed with total RNA extracted from UK1_O using a 5′RACE System kit (Sigma Aldrich), according to the instruction provided by the manufacturer. Gene-specific RT primer (SP1) and PCR primer (SP2) were designed as recommended. The 5’-RACE product was used as a template for a nested PCR reaction using a second primer (SP3). The sequences of primers used in the RACE analysis are shown in Table S2. PCR products were cloned in pGEM-T vector and were sequenced.

### Mutagenesis of *CDR20291_0685* upstream region

Quick Change Lightning Site-Directed Mutagenesis Kit (Agilent Technologies) was used to carry out site-directed mutagenesis whereby the Left Inverted Repeat (LIR) TAGTTGTAAAAGGGTT that flanked the DNA region that undergoes inversion was converted to TACACATGCGAGGGTT. The mutagenic oligonucleotide primers used are listed in Table S2.

### Construction of reporter plasmids and beta-glucuronidase assay

The 1.5 kb upstream DNA regions of *CDR20291_0685* were amplified by PCR using specific primers with *KpnI* and *SacI* (Table S2) recognition sequences. UK1_T and UK1_O chromosomal DNA were used as a template to amplify the upstream region in respective orientations. The upstream region was cloned in the pRPF185 shuttle vector using standard cloning procedures. Plasmid pRPF185 carries a *gusA* gene for beta-glucuronidase under the tetracycline-inducible (*tet*) promoter (Fagan and Fairweather, 2011). The *tet* promoter was removed using *KpnI* and *SacI* digestion and was replaced with the *CDR20291_0685* upstream region to create plasmids pBA042 and pBA045 (Table S1). The inverted repeats in the upstream region were mutagenized as described earlier to create plasmid pBA043 containing the upstream region locked in translucent orientation and plasmid pBA046 containing the upstream region locked in opaque orientation. The control plasmid pBA040 with promoter-less *gusA* was created by digesting with *KpnI* and *SacI* to remove *tet* promoter and then self-ligated after creating blunt ends. Plasmids were introduced into *C. difficile* strains through conjugation. The transconjugants were grown overnight in TY medium in the presence of thiamphenicol (15 µg/ml). Overnight cultures were used as inoculum at a 1:100 dilution to start a new culture. Bacterial cultures were harvested at different time points of growth, and the amount of beta-glucuronidase activity was assessed as described elsewhere (Mani *et al*., 2002). Briefly, the cells were washed and resuspended in 1 ml of Z-buffer (60 mM Na_2_HPO_4_.7H_2_0 pH 7.0, 40 mM NaH_2_PO_4_.H_2_O, 10 mM KCl, 1mM MgSO_4_.7H_2_O, and 50mM 2ME), and lysed by homogenization. The lysate was mixed with 160 µl of 6mM p-nitrophenyl β-D-glucuronide (Sigma) and incubated at 37°C. The reaction was stopped by the addition of 0.4 ml of 1.0 M NaCO_3_. β-Glucuronidase activity was calculated as described earlier (Dupuy and Sonenshein, 1998; Mani *et al*., 2002).

### Electrophoretic mobility shift assay (EMSAs)

For the CodY binding experiment, the upstream region of the *pdcB* with the predicted CodY binding sequence 5’ CATAGATAATTTTTAGAAAAGTATCTAAATTTTCAATAAATAGTAAC 3’ was synthesized. It was labeled with [γ-32 P] dATP-6000 Ci/mmol (PerkinElmer Life Sciences) using T4 polynucleotide kinase. It was then annealed with the complementary oligo to generate a double-stranded DNA probe. The DNA-protein binding reactions were carried out at room temperature for 30 min in 10μl volume containing 1x binding buffer [10mM Tris pH 7.5, 50mM KCl, 50μg BSA, 0.05% NP40, 10% Glycerol, 10 mM GTP and 2mM ILV (Isoleucine, Leucine and, Valine), 100 μg/ml poly dI-dC and 800nM of DNA probe with varying concentration of purified CodY protein. DNA probe in reaction buffer was incubated for 10 min at RT before adding purified CodY-6His protein. The reaction was stopped by adding 5µl of gel loading buffer and electrophoresed at 100V for 1.5 h using 6% 1XTBE gel in 0.5X TBE buffer containing 10 mM ILV. Gels were then dried, and the autoradiography was performed with Molecular Dynamics Phosphor-Imager technology.

### Construction and complementation of *C. difficile* UK1::*pdcB* mutant strains

The UK1::*pdcB* mutant in the UK1 strain was created using the ClosTron gene knockout system as described previously (Heap *et al*., 2010). Briefly, for *pdcB* disruption, the group II intron insertion site between nucleotides 840 and 841 in the *pdcB* gene in the sense orientation was selected using a web-based design tool called the Perutka algorithm. The designed retargeted intron was cloned into pMTL007-CE5 as described previously (Girinathan *et al*., 2016; Govind and Dupuy, 2012). The resulting plasmid pMTL007-CE5::*pdcB*-840-841s was transferred into *C. difficile* UK1 cells by conjugation. The potential Ll.ltrB insertions within the target genes in the *C. difficile* chromosome were conferred by the selection of lincomycin-resistant transconjugants in 20 μg/ ml lincomycin plates. PCR using gene-specific primers (Table S2) in combination with the EBS-U universal was performed to identify putative *C. difficile* mutants. *C. difficile pdcB* mutants were complemented by introducing pBA048, which contains the PdcB*-*EAL domain, and pBA050, which contains the PdcA-EAL domain under inducible promoter, through conjugation.

### Cyclic-di-GMP measurement

c-di-GMP was measured as described previously (Girinathan *et al*., 2017), with few modifications. Briefly, *C. difficile* strains were grown in 50 ml TY medium for 16 hours. OD_600_ was recorded and dilution plating was carried out to determine the number of CFU. Cells were centrifuged and pellets were washed once with TE buffer (10 mM Tris [pH 7.5], 1 mM EDTA, pH 8). Pellets were resuspended in extraction solvent consisting of methanol (MeOH): acetonitrile: distilled water (dH_2_O) in 40:40:20 ratio plus 0.1 N formic acid. The mixture was incubated in -20°C for 30 min. The extract was harvested by centrifugation. Samples were sent to Kansas University Biochemistry Core facility to be analyzed by high-pressure liquid chromatography (HPLC)-tandem mass spectrometry (MS/MS). To determine the amount of intracellular c-di-GMP, a standard curve was first obtained by analyzing the serial dilution of pure c-di-GMP (Sigma-Aldrich). The peak intensity of each sample was fitted to the standard curve and the value was divided by the total intracellular volume of bacteria extracted. To determine the total intracellular volume of bacteria, the volume of one cell was multiplied by the number of cells extracted, which was based on CFU counts. The volume of one cell was estimated as a cylinder and was determined by measuring the radius and length of the bacterial cell from the electron micrographs.

## Data Sharing statement

Data sharing is not applicable to this article as no new data were created or analyzed in this study.

## Figure Legends

**Figure S1. Swimming motility and toxin production of UK1_T and UK1_O strains. A**. Swimming motility exhibited by UK1_T and UK1_O in TY+0.5% agar. **B**. Cytoplasmic toxin levels produced by UK1_T and UK1_O morphotypes during the stationary phase of growth. Three biological replicates were used to carry out the experiments, and data were analyzed by a two-tailed unpaired *t*-test with Welch’s correction (NS-Not Significant).

**Figure S2. UK1_O undergoes DNA inversion upstream of *CDR20291_0685*. A**. NCBI BLASTn result showing the perfect alignment of 168 bp segment of the upstream region of *CDR20291_0685* from UK1_T with the published R20291 sequence. **B**. NCBI BLASTn result showing the mismatch alignment of the 168 bp segment of the upstream region of *CDR20291_0685* from UK1_O strain with the published R20291 sequence. The invertible region is flanked by inverted repeats shown by the black lines.

**Figure S3. Construction and confirmation of the *pdcB* mutation in *C. difficile* UK1 strain. A**. Schematic representation of ClostTron (group II intron) mediated insertional inactivation of *pdcB* gene in *C. difficile*. **B**. PCR verification of the intron insertion in *pdcB in* UK1 strain, conducted with intron-specific primer EBS universal [EBS(U)] with *pdcB* specific primers ORG 771 and ORG 772.

**Figure S4. Increased c-di-GMP is associated with increased swarming motility phenotype in UK1_T strain. A**. Representative swarming motility plates of UK1_T, UK1_O, UK1::*pdcB* mutant, UK1::*pdcB* + PdcB-EAL and UK1::*pdcB* + PdcA-EAL complemented strains after 72 hours of incubation at 37°C.

**Figure S5. Orientation of *pdcB* and *cmrRST* upstreams in UK_T and UK_O cells. A**.PCR amplification of the upstream region of *pdcB* using primer mixture containing primers ORG886/887/888. Amplification with ORG886/888 results in a product size of 325 bp, while amplification with ORG 887/888 results in a product size of 215 bp. **B**. PCR amplification of the upstream region of *cmrRST* using primer mixture containing primers ORG879/880/881. Amplification with ORG879/880 results in a product size of 410 bp, while ORG 880/881 results in 540 bp PCR product.

