## Supplemental Material for "Phase variable expression of *pdcB*, a phosphodiesterase influences sporulation in *Clostridioides difficile*"

Table S1

| **Bacterial strain or plasmid** | **Relevant features or genotype** | **Reference** |
|---|---|---|
| *Clostridioides difficile* UK1 | Clinical isolate | (Sorg and Sonenshein, 2010) |
| *Escherichia coli* DH5α | *endA1 recA1 deoR hsdR17 ($r_K^-$ $m_K^+$)* | NEB |
| *Escherichia coli* S17-1 | Strain with integrated RP4 conjugation transfer function; favors conjugation between *E. coli* and *C. difficile* | (Teng et al., 1998) |
| *Clostridioides difficile UK1::pdcB* | UK1 with intron insertion within *pdcB* (*CDR20291_0685*) | This study |
| *Clostridioides difficile* UK1::*codY* | UK1 with intron insertion within *codY* | (Nawrocki et al., 2016) |
| pMTL007-CE5 | ClosTron plasmid | (Heap et al., 2010) |
| pMTL007-CE5:Cdi-*pdcB*-840-841s | pMTL007-CE5 with group II intron targeted to *pdcB* | This study |
| pRPF185 | *E. coli*/ *C. difficile* shuttle plasmid | (Fagan and Fairweather, 2011) |
| pBA042 | pRPF185 containing 1.5 kb upstream region of *pdcB* in Translucent orientation. | This study |
| pBA043 | pRPF185 containing ~1.5 kb *pdcB* upstream region locked in Translucent orientation | This study |
| pBA045 | pRPF185 containing 1.5upstream region of *pdcB* in Opaque orientation. | This study |
| pBA046 | pRPF185 containing ~1.5 kb *pdcB* upstream region locked in Opaque orientation | This study |
| pBA048 | pRPF185 containing PdcB-EAL domain | This study |
| pBA050 | pRPF185 containing PdcA-EAL domain | This study |
| pBA051 | pRPF185 containing the *tet* promoter alone | This study |
| pBA052 | pRPF185 containing *dccA* under *tet* promoter | This study |

#### Table S2 Oligonucleotides used for PCR reactions

| Name | Sequence (5' → 3') | Description |
|---|---|---|
| EBS-U | CGAAATTAGAAACTTGCGTTCAGT AAAC | Group II intron specific primer |
| ORG771 | GACTGAGCTCTGGGGAAATGTGTTTGATGA GATAAAAAAATTAGTAAAATA | Forward primer with with SacI to clone *CDR20291_0685* (*pdcB*) in pRPF185 |
| ORG772 | AACTGGATCCTCACTTATCGTCGTCATCCT TGTAATCCTTTGATAGTTGAAATTTATAAA ATTCAGAGGCAC | Reverse primer with with BamHI to clone *CDR20291_0685* (*pdcB*) in pRPF185 |
| ORG788 | GAGCTCAAGAGGAGTGGTTGAAAATGGCAA CAAGACCTATAGAAATAG | Forward primer with SacI to amplify *recV* |
| ORG789 | GGATCCTTAATGATGATGATGATGATGACC AATAAAGAAATTTTCACTAGCTTCATTAAT AGCTTGCTGAG | Reverse primer with BamHI and 6X His to amplify *recV* |
| ORG844 | GTTGTAAAAAAGTTACTATTTATTGAAAAT TTAGATACTTTTCTAAAATTATCTATG | Forward primer for CodY binding site in *pdcB* upstream region in Opaque orientation |
| ORG845 | CATAGATAATTTTAGAAAAGTATCTAAATT TTCAATAAATAGTAACTTTTTTACAAC | Reverse primer for CodY binding site in *pdcB* upstream region in Opaque orientation |
| ORG852 | GGTACCAGTTTAGGATAAAGTATTGCAAGA ACCAATCAG | Forward primer with KpnI site to clone ~1.5 kb upstream region of *pdcB* gene |
| ORG853 | GAGCTCCTTTTCCCCCTACAATATTACTAT TAGTGTAGTTTAATCAAC | Reverse primer with SacI site to clone ~1.5 kb upstream region of *pdcB* gene |
| ORG866 | ATGTATATTTTTATAGCTATTTTATTATAA AAAAAAAGAACCCTCGCATGTGTAAGGGTT ACTTTTTTTCTCTATTTTTTTTATTATATC ACTATTTTTTCCCTATTTCAATATTTTGAC ATTTTTTCCATTATACTGA | Forward primer to mutagenize inverted repeat from translucent orientation |
| ORG867 | TCAGTATAATGGAAAAAATGTCAAAATATT GAAATAGGGAAAAAATAGTGATATAATAAA AAAAATAGAGAAAAAAAGTAACCCTTACAC ATGCGAGGGTTCTTTTTTTTTATAATAAAA TAGCTATAAAAATATACAT | Reverse primer to mutagenize inverted repeat from translucent orientation |
| ORG868 | CATAAATTTAACTATTAACATAGATAATTT TAGAAAAGTATCTAAATTTTCAATAAATAG TAACTTTCGCATGTGTAAGGGTTACTTTTT TTCTCTATTTTTTTTATTATATCACTATTT TTTCCCTATTTCAATATTTTGAC | Forward primer to mutagenize inverted repeat from opaque orientation |
| ORG869 | GTCAAAATATTGAAATAGGGAAAAAATAGT GATATAATAAAAAAAATAGAGAAAAAAAGT AACCCTTACACATGCGAAAGTTACTATTTA TTGAAAATTTAGATACTTTTCTAAAATTAT CTATGTTAATAGTTAAATTTATG | Reverse primer to mutagenize inverted repeat from opaque orientation |
| ORG879 | GTATTATTTTGGTAAATATATTGTTACAAA AGGTTTATATTTTGC | Forward primer (*cmrRST* upstream): specific for orientation similar to published (410 bps) |

| | | |
|---|---|---|
| ORG880 | GGAAATATTGACAAAATAATATTACAATGTTAGAATAA | Reverse primer common for both orientation or *cmrRST.* |
| ORG881 | AGTATAATGCTATTATAATAAGAAAATAACTTTTTTATAAACATTGAGAT | Forward primer (*cmrRST* upstream): specific for flipped orientation (540 bps) |
| ORG882 | GAGCTCGGGGGAAAAGATGTATCCAGAAGATGGAGATAATTATTTAGATTTATTTAAACA | Forward primer with *SacI* to clone PdcB-EAL in pRPF185 |
| ORG883 | GGATCCTCACTTTGATAGTTGAAATTTATAAAATTCAGAGGCACTTACAGGTCTTCC | Reverse primer with *BamHI* to clone PdcB-EAL in pRPF185 |
| ORG884 | GAGCTCGGAGGAGATAAGATGCAAGAAATATTAAAAAATAAAATGTATTCAT | Forward primer with *SacI* to clone PdcA-EAL in pRPF185 |
| ORG885 | GGATCCTTAATTATCTAGCTTTAAAAGGTCAAAGATTTCTGTTGCTGTCTGCGGTTTTCC | Reverse primer with *BamHI* to clone PdcA-EAL in pRPF185 |
| ORG886 | TAATAAAATAGCTATAAAAATATACATATCAAATCAAATTAAG | Forward primer to amplify the flip orientation of *pdcB* upstream in Translucent strain. |
| ORG887 | CTTCTTAATTTGATTTGATATGTATATTTTTATAGC | Forward primer to amplify the flip orientation of *pdcB* upstream in Opaque strain. |
| ORG888 | CTGGATTTTTAAATTTATGTACTTTAAAATGGATATCC | Reverse primer to amplify the flip orientation both translucent and opaque of *pdcB* upstream. |
| ORG846 | GGTCTAAACTTAAAGAGTTCGATTCTTCTCTCATCTTTTCCCCCTACAATATTACTATTAGTG | SP1 primer for 5' RACE |
| ORG847 | CTCTAAGTATTTCATTAAAAAGATGTTAGTAATGACCACTAAATAAAAGTGGACGATTTTGTGG | SP2 primer for 5' RACE |
| ORG848 | CAATTATTTTATCCATGTAAAAATCATGTACTTTTATCTCAACGAATGTCCC | SP3 primer for 5' RACE |
| ORG921 | ATGGAAACTATGTACGTAATAATTAAAATAAAA | Forward primer to amplify *pdcB* promoter_1 |
| ORG922 | ACATTGACATTCTAATTCTGTAGTGTAATCGTA | Forward primer to amplify *pdcB* promoter_2 |
| ORG925 | ACTATTTATTGAAAATTTAGATACTTTTCTAAAATTATCTATG | Forward primer to amplify *pdcB* promoter_3 |
| ORG926 | ATACTTTTCTAAAATTATCTATGTTAATAGTTAAAT | Forward primer to amplify *pdcB* promoter_3 |

Supplementary Figure 1

A

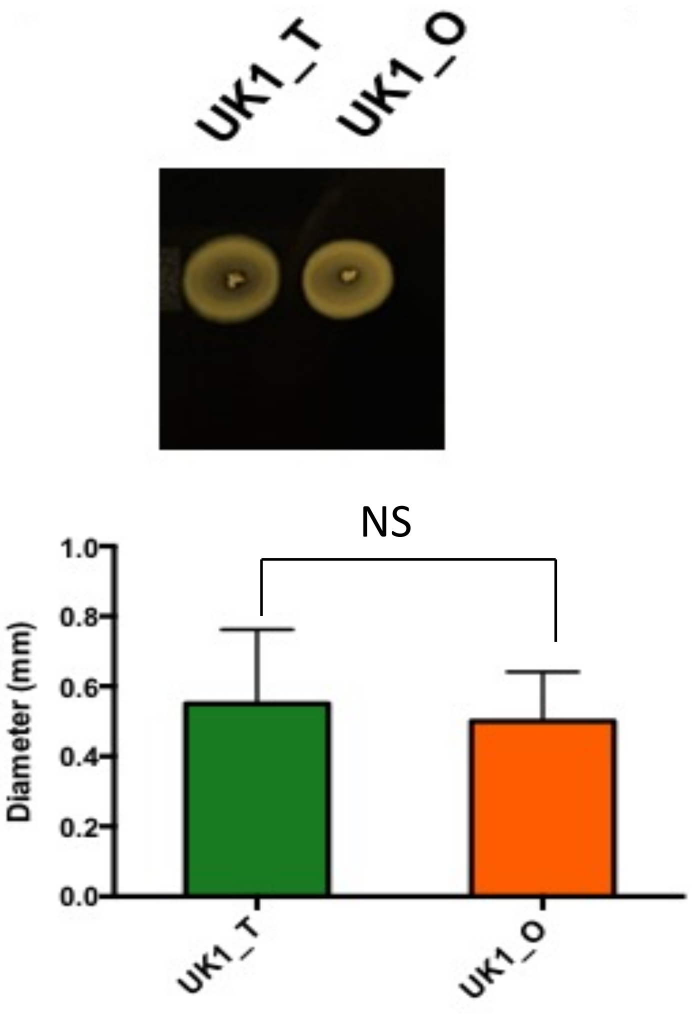


B

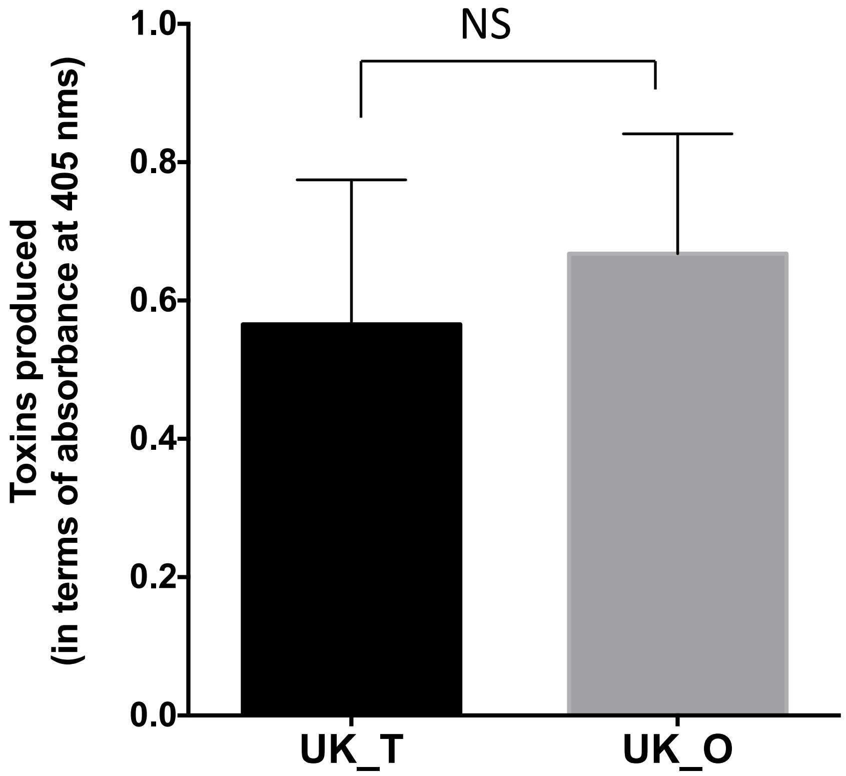

### Supplementary Figure 2

**A**

```
1       AATGTCaaaatattgaaatagggaaaaaatagtgatataataaaaaaaatagagaaaaaa  60      UK_T
862733  AATGTCAAAATATTGAAATAGGGAAAAAATAGTGATATAATAAAAAAAATAGAGAAAAAA  862792  R20291 Published

61      aGTAACCCTTAGTTGTAAAAGGGTTCtttttttttATAATAAAATAGCTATAAAAATATA  120     UK_T
862793  AGTAACCCTTAGTTGTAAAAGGGTTCTTTTTTTTTATAATAAAATAGCTATAAAAATATA  862852  R20291 Published

121     CATATCAAATCAAATTAAGAAGTATTTCATTTCTAAGAAATATCCTAACATaaaaacaaa  180     UK_T
862853  CATATCAAATCAAATTAAGAAGTATTTCATTTCTAAGAAATATCCTAACATAAAAACAAA  862912  R20291 Published

181     aaaTGTTCATAAATTTAACTATTAACATAGATAATTTTTAGAAAAGTATCTAAATTTTCA  240     UK_T
862913  AAATGTTCATAAATTTAACTATTAACATAGATAATTTTTAGAAAAGTATCTAAATTTTCA  862972  R20291 Published

241     ATAAATAGTAACTTTTTTACAACAAATGGAAACTATGTACGTAATAATTAAAATAAAAAA  300     UK_T
862973  ATAAATAGTAACTTTTTTACAACAAATGGAAACTATGTACGTAATAATTAAAATAAAAAA  863032  R20291 Published

301     TATTATTTTATATTTATAAACATTGA  326     UK_T
863033  TATTATTTTATATTTATAAACATTGA  863058  R20291 Published
```

**B**

```
1       AATGTCaaaatattgaaatagggaaaaaatagtgatataataaaaaaaatagagaaaaaa  60      UK_O
862733  AATGTCAAAATATTGAAATAGGGAAAAAATAGTGATATAATAAAAAAAATAGAGAAAAAA  862792  R20291 Published

61      agtaacccttagttgtaaaaaaGTTACTATTTATTGAAAATTTAGATACTTTTCTAAAAT  120     UK_O
862793  AGTAACCCTTAGTTGTAAAAGGGTT-CTTTTTTTTTATAAT-AAAATA--GCTATAAAAA  862848  R20291 Published

121     TATCTATGTTAATAGTTAAATTTATGAACAttttttgtttttATGTTAGGATAT--T-TC  177     UK_O
862849  TATACATATCAA-A-TCAAA-TTAAGAA-GTATTTCATTTCTAAG--A-AATATCCTAAC  862901  R20291 Published

178     TTAGAAATGAAATA-CTTC-TTAATTTGA-T-TTGATATGTAT-ATTTTT--ATAGCTAT  230     UK_O
862902  ATAAAAACAAAAAATGTTCATAAATTTAACTATTAACATAGATAATTTTTAGAAAAGTAT  862961  R20291 Published

231     -TTTATTATaaaaaaaaaG-AACCCTTTTACAACAAATGGAAACTATGTACGTAATAATT  288     UK_O
862962  CTAAATTTTCAATAAATAGTAACTTTTTTACAACAAATGGAAACTATGTACGTAATAATT  863021  R20291 Published

289     AAAATAAAAAATATTATTTTATATTTATAAACATTGA  325     UK_O
863022  AAAATAAAAAATATTATTTTATATTTATAAACATTGA  863058  R20291 Published
```

Supplementary Figure 3

A.

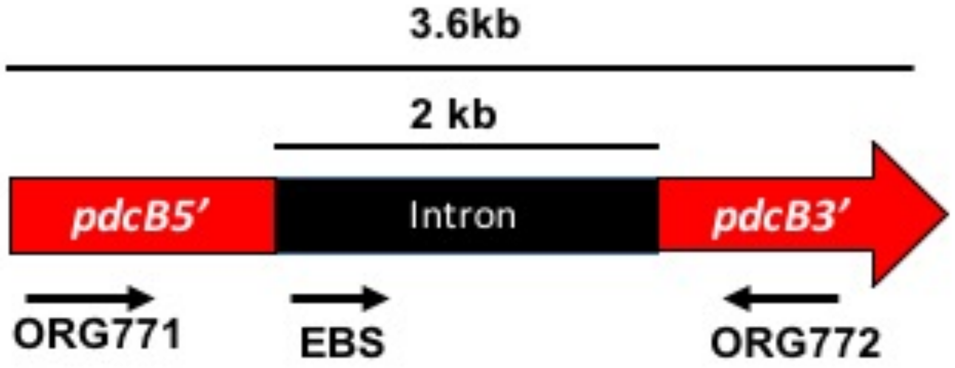


B.

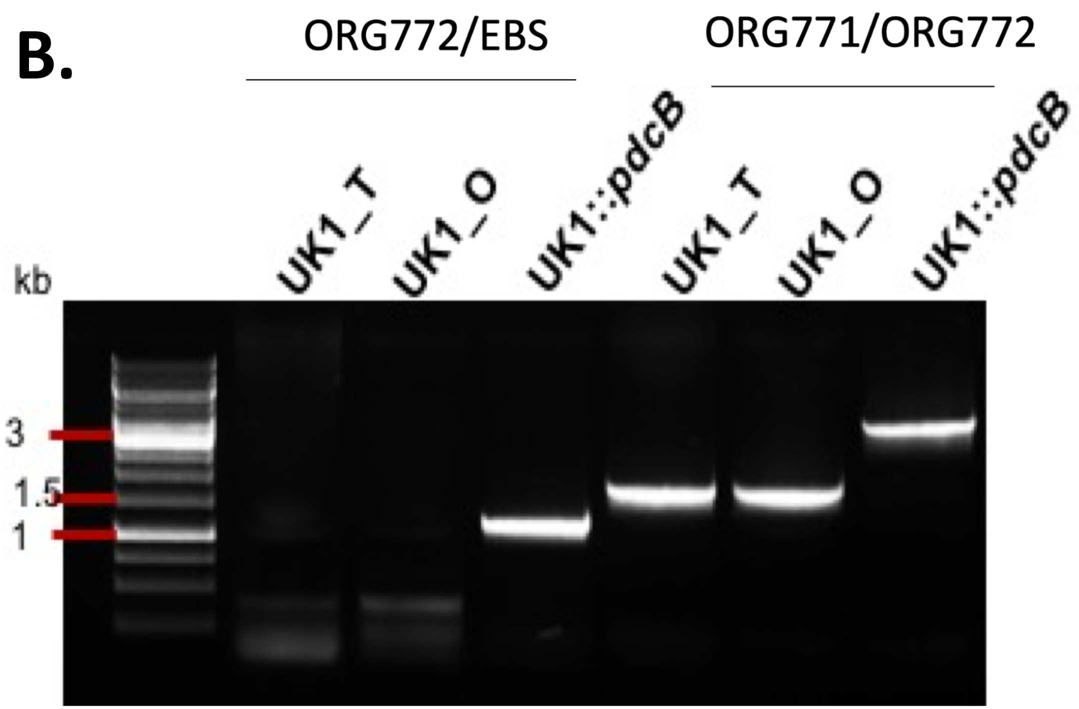

### Supplementary Figure 4

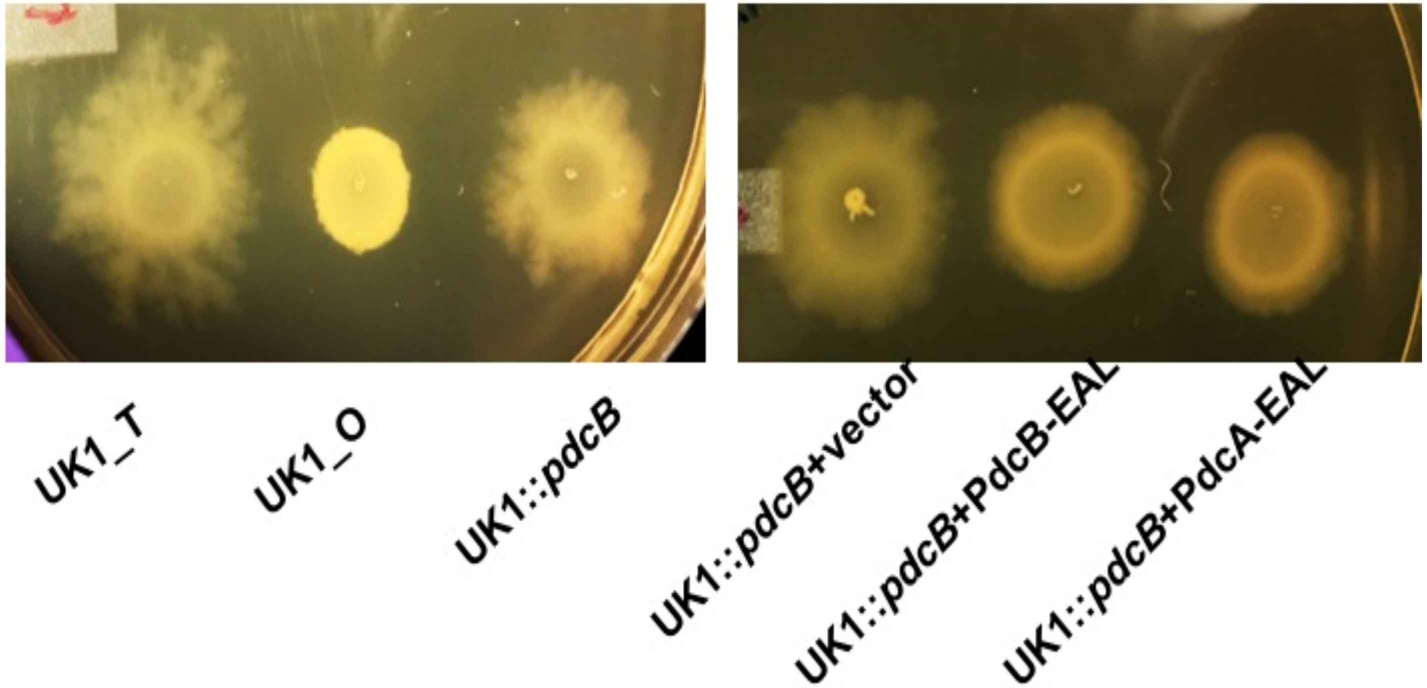

### Supplementary Figure 5

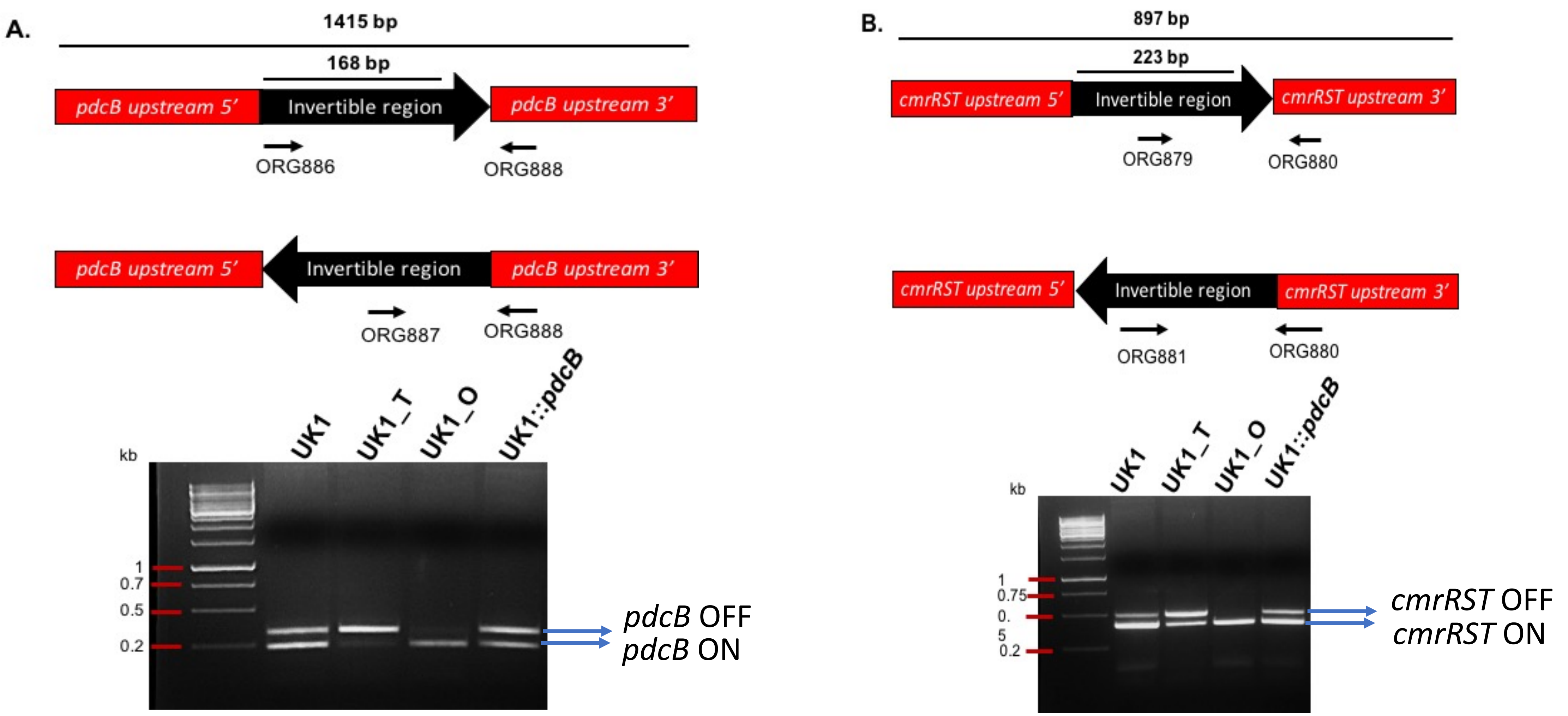
